# Research article, Biological Sciences - Anthropology, Genetics Of cats and men: the paleogenetic history of the dispersal of cats in the ancient world

**DOI:** 10.1101/080028

**Authors:** Claudio Ottoni, Wim Van Neer, Bea De Cupere, Julien Daligault, Silvia Guimaraes, Joris Peters, Nikolai Spassov, Mary E. Prendergast, Nicole Boivin, Arturo Morales, Adrian Bălăşescu, Cornelia Becker, Norbert Benecke, Adina Boronenanţ, Hijlke Buitenhuis, Jwana Chahoud, Alison Crowther, Laura Llorente, Nina Manaseryan, Hervé Monchot, Vedat Onar, Marta Osypińska, Olivier Putelat, Jacqueline Studer, Ursula Wierer, Ronny Decorte, Thierry Grange, Eva-Maria Geigl

## Abstract

The origin and dispersal of the domestic cat remain elusive despite its importance to human societies around the world. Archaeological evidence for domestication centers in the Near East and in Egypt is contested, and genetic data on modern cats show that *Felis silvestris lybica*, the subspecies of wild cat inhabiting at present the Near East and Northern Africa, is the only ancestor of the domestic cat. Here we provide the first broad geographic and chronological dataset of ancient cat mtDNA sequences, drawing on archaeological specimens from across western Eurasia and northern and eastern Africa, dating from throughout the Holocene and spanning ~9,000 years. We characterized the ancient phylogeography of *F. s. lybica,* showing that it expanded up to southeastern Europe prior to the Neolithic, and reconstructed the subsequent movements that profoundly transformed its distribution and shaped its early cultural history. We found that maternal lineages from both the Near East and Egypt contributed to the gene pool of the domestic cat at different historical times, with the Near Eastern population providing the first major contribution during the Neolithic and the Egyptian cat spreading efficiently across the Old World during the Classical period. This expansion pattern and range suggest dispersal along maritime and terrestrial routes of trade and connectivity. Late trait selection is suggested by the first occurrence in our dataset of the major allele for blotched-tabby body marking not earlier than during the Late Middle Ages.

**Significance:** The cat has long been important to human societies as a pest-control agent, object of symbolic value, and companion animal, but little is known about its domestication process and early anthropogenic dispersal. Our DNA analyses of geographically and temporally widespread archaeological cat remains show that while the cat’s world-wide conquest began in prehistoric times, when tamed cats accompanied humans on their journeys over land and sea, it gained momentum during the Classical period, when the Egyptian cat successfully spread throughout the ancient world. The appearance of a new coat pattern at the end of the Middle Ages suggests late breeding control that might explain the semi-domestic status of the cat. This distinguishes the domestication process of cats from that of most other domesticates.

## Introduction

The domestic cat is present on all continents and in the most remote regions of the world, and its evolutionary success is unquestioned. While nowadays it is among the most cherished companion animals in the Western world, for ancient societies barn cats, village cats and ship’s cats provided critical protection against vermin, especially rodent pests responsible for economic loss and disease (1). Archaeological and historical evidence suggest that the global spread of domestic cats was shaped by their close association with humans; accordingly, studying their dispersal has the potential to reveal aspects of human history such as distant trade relations including ancient land and maritime trade routes and to contribute to a better understanding of how humans have reshaped global biodiversity through introduced animals (e.g., (2–4)). Given their low frequency in the archaeological record, patterns of cat domestication and translocation remain largely elusive, explaining why cats have been less studied than other domesticates, such as various species of livestock or dogs, particularly from a genetic perspective. Wildcats (*Felis silvestris*) are distributed all over the Old World. Current taxonomy distinguishes four wild subspecies, *F. s. silvestris, F. s. lybica, F. s. ornata and F. s. cafra*, geographically partitioned across the Old World (5). Only one of them, the North African/Southwest Asian *F. s. lybica,* was ultimately domesticated (6). Wildcats are solitary, territorial hunters and lack a hierarchical social structure (7, 8), features that make them poor candidates for domestication (9). Indeed, zooarchaeological evidence points to a commensal relationship between cats and humans lasting thousands of years before humans exerted substantial influence on their breeding (10–12). Throughout this period of commensal interaction, tamed and domestic cats became feral and/or intermixed with wild *F. s. lybica* or other wild subspecies once introduced into their native range, as is common today (13). This process most likely caused high levels of gene flow between wild and domestic populations that lasted even longer than reported for other domesticates (14–16). Accordingly, the domestication process has seemingly not profoundly altered the morphological, physiological, behavioral and ecological features of cats (12), in contrast to what has been observed, for example, for dogs (17). This limits the potential for morphological discrimination of wild and domestic cats for most archaeological specimens, particularly for the African wildcat, for which no extensive morphological and osteometric studies have hitherto been performed (10).

Difficulties in tackling cat domestication are also linked to a paucity of cat remains in the zooarchaeological record (apart from abundant Egyptian mummies of the Greco-Roman period), as cats were not a subsistence species and are therefore rarely found in refuse contexts. As a consequence, current hypotheses about early cat domestication rely on only a few zooarchaeological case studies. A complete skeleton found in Cyprus in association with a human burial dated to ca. 7,500 BCE suggests that cats were likely tamed by early Neolithic sedentary communities growing cereals, concomitant with the emergence of commensal rodents (11). Similarly, the skeletons of six cats in an elite Predynastic cemetery in Egypt, ca. 3,700 BCE, suggest a close cat-human relationship in early ancient Egypt (10). The iconography of Pharaonic Egypt constitutes a key source of information about the species’ relationship with humans, and has motivated the traditional belief that cat domestication took place in Egypt (1, 18). Animals in Egyptian art are usually accurately portrayed and the numerous depictions from the 2^nd^ millennium BCE document a progressive tightening of the relationship between human and cat, as illustrated in particular by the popularization of the motif of the “cat under the chair” of seated women after ca. 1,500 BCE (1, 18). Thus, ancient societies in both the Near East and Egypt could have played key roles in cat domestication, but it is not known to what extent these early tamed cats contributed to modern domestic cats, nor are the modalities of their dispersal across the Old World understood.

Modern genetic data have also been unable to resolve these questions. Analyses of nuclear short tandem repeats (STR) and 16% of the mitochondrial DNA (mtDNA) genome in extant wild and domestic cats (6) have identified five geographically distinct clades (I to V, Fig. S1) representing the four *Felis silvestris* subspecies as well as *Felis bieti* (Chinese mountain cat), and revealed that modern domestic cats share clade IV with *F. s. lybica* and must therefore originate from this subspecies. The modern domestic cat mtDNA pool was traced back to five deeply divergent subclades (IV-A to IV-E), representing multiple wildcat lineages incorporated over time and space (7). These subclades lack a phylogeographic structure, which may reflect either poor sampling of the truly wild modern *F. s. lybica*, particularly in its African range, or extensive gene flow between wild and domestic populations following the dispersal of domestic cats. The analysis of full genomes revealed a low level of differentiation between modern wild and domestic cats, presumably due to prolonged hybridization (19), but could not address questions concerning the chronology and geography of cat domestication and dispersal.

Due to a lack of temporal depth, drawing on modern genetic data alone to explore domestication is open to pitfalls (20), especially as ancient DNA (aDNA) studies demonstrate that past demographic events can alter phylogeographic structure (21). Ancient DNA analysis has the power to explore whether a fine phylogeographic structure existed prior to the domestication of *F. s. lybica* and whether, when, and how it was reconfigured over time in response to human intervention. In this way, the phylogeography of ancient cats has the potential to document the cat domestication process. Ancient DNA can also be used to monitor changes in the frequencies of alleles underlying phenotypic traits that differentiate wild and domestic populations (22). Indeed, such changes reveal human-mediated selection of traits, reflecting the extent of the breeding control exerted on animal populations during the domestication process. In particular, variations in coat pattern have been among the first traits to be associated with the domestic status of some animals, such as horses (22).

To address questions related to the contribution of the two purported centers of cat domestication, the Near East and Egypt, and the history of human-mediated dispersal and trait selection in the domestic cat, we analyzed ancient cat remains from Europe, North and East Africa, and Southwest Asia (SWA), spanning around 9,000 years, from the Mesolithic to the 19^th^ century CE. We investigated maternal lineages to trace the evolution of the ancient phylogeographic structure of *F. s. lybica,* as well as a genetically defined coat color marker, the blotched tabby marking (23), to monitor a phenotypic change reflecting human-driven selection along the domestication pathway.

## Results

### Strategy for data acquisition

In order to screen and analyze in parallel a large number of ancient samples, many of which were expected to be poorly preserved due to higher temperature and sometimes tropical burial environments, we applied the “aMPlex Torrent approach” that combines the sensitivity of multiplex PCR with the power and throughput of next-generation sequencing (24). We targeted 42 informative SNPs in nine short regions distributed across the mitochondrial ND5, ND6 and CytB genes that recapitulate the most salient features of the phylogeny obtained with these complete genes (Fig. S1). We obtained mtDNA sequences from 209 out of 351 ancient samples (59%; Dataset S1, S2), with expectedly lower success rates for old samples from hot environments (Fig. S2).

The mtDNA phylogeny (Fig. 1b) reconstructed from 286 bp sequenced in our ancient samples alongside modern data from the literature (6) clearly separates the four clades of *Felis silvestris* and *Felis bieti* (posterior probabilities >0.88, Fig. S3, SI Methods) and the five subclades of *F. s. lybica* (posterior probabilities >0.77). We examined the phylogeographic pattern and its changes across time by grouping the mtDNA haplotypes from our study into nine time bins (Fig. 1c).

**Figure 1.**
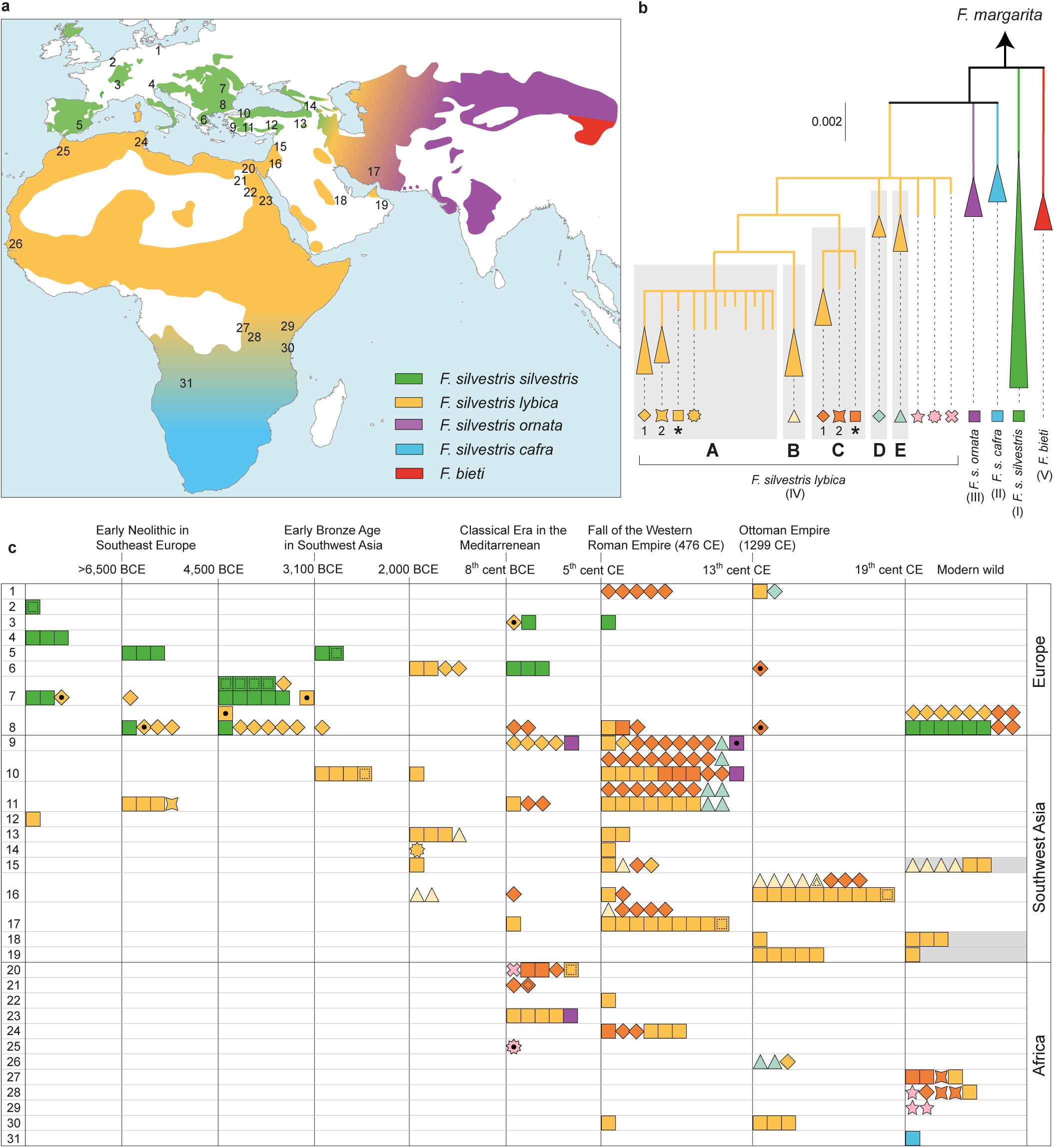
Spatiotemporal representation of cat maternal genealogies. (**a**) Map showing the present-day distribution of *Felis silvestris* (5) with the geographic range of each subspecies as reported in literature (6). (**b**) Tree of mtDNA lineages observed in our ancient samples and in modern wild and domestic cats from literature (6). (**c**) Spatiotemporal depiction of ancient cat haplotypes as depicted with symbols from the tree (b). Rows represent the approximate geographic provenance of the samples as reported in the map (a) whereas the columns pertain to chronological periods. A dot inside the symbols indicates AMS radiocarbon dated samples; dashed lines inside the symbols indicate incomplete mtDNA profiles; Near Eastern modern wildcats from literature (6) are indicated by grey shaded bins. Numbers in (a) and in (c) represent the approximate geographic locations of the sites from which the samples are derived as follows: **Germany**: #1) Berlin, Ralswiek. **Belgium**: #2) Trou Chaleux. **France**: #3) Entzheim-Geispolsheim, Entzheim-Lidl, Illfurth Buergelen. **Italy**: #4) Galgenbühel/Dos de la Forca. **Spain**: #5) Cova Fosca, Tabernas, Valencina. **Greece**: #6) Kastanas, Kassope, Tiryns. **Romania**: #7) Cheia, Hârşova, Icoana, Pietrele, Vităneşti. **Bulgaria**: #8) Arbanas, Budjaka, Dolnoslav, Durankulak, Garvan, Hotnitsa, Jambol, Kabile, Koprivec, Sliven. **Turkey**: #9) Didyma, Kadıkalesi, #10) Demircihüyük, Yenikapı, #11) Sagalassos, Bademağacı, #12) Aşıklı Höyük, #13) Norşun Tepe, Hassek Höyük, Lidar Höyük, Anzaf. **Armenia**: #14) Keti, Dvin. **Syria**: #15) Tell Guftan, Shheil 1, Qasr al-Hayr al-Sharqi. **Lebanon**: #15) Sidon. **Jordan**: #16) Aqaba, Tall al-'Umay'ri, Hesban, Khirbet es-Samra. **Iran**: #17) Siraf. **Saudi Arabia**: #18) Al-Yamama. **Oman**: #19) Qalhât. **Egypt**: #20) mummies from Natural History Museum and British Museum (London, UK), #21) Oxyrhynchus, #22) Shenhur, #23) Berenike. **Tunisia**: #24) Bigua. **Morocco**: #25) Mogador. **Senegal**: #26) Gorée. **Congo**: #27) modern wildcats. **Burundi-Rwanda**: #28) modern wildcats. **Kenya**: #29) modern wildcats. **Tanzania**: #30) Songo Mnara, Unguja Ukuu, Mapangani Cave. **Angola**: #31) modern wildcats.

### Ancient European wildcats

Covering more than nine millennia, we found the mtDNA clade I, representative of European wildcats (*F. s. silvestris*), exclusively in Europe. From the Mesolithic to the 8^th^ century BCE in Western Europe (geographic locations #1-5 in Fig. 1a,c), all cats analyzed (9/9) carried clade I mtDNA, whereas in Southeast Europe (#6-8) we observed similar frequencies of clade I (N=13, 42%) and clade IV (N=18, 58%), representative of *F. s. lybica*. The latter was mostly represented by one of the lineages of subclade IV-A, named here IV-A1 (Fig. S1, S3), the earliest occurrence of which, in our dataset, dates back to 7,700 BCE in Romania (#7) (Dataset S1), and which is still present today in European wild (#8) and domestic cats (6).

### Anatolian cats from the Neolithic to the Bronze Age

A mitotype belonging to subclade IV-A (named here IV-A*, see Methods section) was predominant (12/14) from ca. 8,000 to the 8^th^ century BCE in Anatolia (#10-13) (Fig. 1a-c). Its range may have also extended to Lebanon (#15). The frequencies of IV-A1 and IV-A* found in Southeast Europe and Anatolia, respectively, are significantly different (Fisher’s exact test; P<0.001), suggesting a phylogeographic structure that mirrors the original distribution of genetically distinct wildcat populations carrying *F. s. lybica* mtDNA. The earliest occurrence of IV-A* outside the Anatolian range in our dataset was detected in two directly radiocarbon-dated specimens from Southeast Europe, in Bulgaria (4,400 BCE) and Romania (3,200 BCE), clearly postdating the introduction of Neolithic farming practices, and in two Late Bronze/Iron Age cats (ca. 1,200 BCE) from Greece.

### Ancient Levant and Africa

Owing to very poor DNA preservation in ancient archaeological contexts in the Levant and northern Africa, we inferred the original distribution of the other subclades (IV-B/E) by taking into account their temporal appearance in our dataset. We found IV-B in three ancient cat remains dated to the 1^st^ millennium BCE from Southeast Anatolia and Jordan (#13, #16, Fig. 1a-c), the 6^th^ century BCE in Syria and later in Jordan (#15-16). This clade is still found in modern wildcats from Israel (6, 25). These data suggest that this subclade was mainly restricted to a Levantine range throughout history. Outside of this range, IV-B was found only in Medieval Iran (#17) at very low frequencies (7%).

In Africa, two lineages of IV-C (named IV-C1 and IV-C*) were detected in five out of seven cats (including three mummies) from Egypt with dates ranging from the 7^th^ century BCE to the 4^th^ century CE (#20-21, Fig. 1a-c). The original range of IV-C may have extended from Egypt along the Nile River as far south as Congo and Burundi (#27-28), where we detected in modern wildcats a novel lineage (IV-C2, Fig. 1) not yet described in the mtDNA pool of present-day domestic cats.

Subclades IV-D and IV-E were found at low frequencies solely in recent temporal bins of our ancient dataset (#1, #9-11), most likely as a result of human-mediated dispersal. Their basal position in clade IV, shared with lineages found in ancient African cats (light pink symbols in Fig. 1 b-c, #20, #25, #28-29) and not detected so far in domestic cats, may suggest an African origin.

### The dispersal of Egyptian cats

Outside Africa, from the 8^th^ century BCE to the 5^th^ century CE, we found IV-C1 in two Late Roman-era cats from Turkey (#11, Fig. 1a-c), in one Classical/Roman-era cat from Jordan (#16), and in two Roman-era cats from Bulgaria (#8). This range expansion is more evident between the 5^th^ and 13^th^ centuries CE, when the two IV-C lineages found in ancient Egyptian cats became significantly more frequent both in Europe (78%; 7/9) and in SWA (50%; 28/56). By contrast, none of the 41 European and 18 Southwest Asian cats from archaeological contexts predating the 8^th^ century BCE possessed IV-C haplotypes (Fisher’s exact test; P<0.001 in both cases).

The territorial behavior of cats and the rapid reconfiguration of the phylogeographic pattern observed in Europe and SWA suggest that cats carrying IV-C haplotypes were spread by humans throughout the eastern Mediterranean by Classical times. Further expansion occurred during the Medieval period, with the IV-C1 haplotype identified at the Viking trading port of Ralswiek on the Baltic Sea (#1, Fig. 1a-c) by the 7^th^ century CE, and the Iranian port of Siraf by the 8^th^ century CE (#17). In the Balkans, IV-C1 persisted throughout Medieval times up to the present (#8). Translocation of cats over even longer distances is witnessed by the presence of Asian wildcat (*F. s. ornata*) mtDNA at the Roman-Egyptian port of Berenike on the Red Sea (1^st^–2^nd^ century CE, #23, Fig. 1a-c) and at Medieval coastal sites in Turkey (#9-10).

### Coat color

To develop a temporal framework of the emergence of a variation in coat color typical of domestic cats we investigated the best-characterized coat color allelic variations involving three single nucleotide polymorphisms (SNPs) in the *Transmembrane Aminopeptidase A* (*Taqpep*) gene controlling the tabby coat pattern (23). We found that the recessive allele responsible for the blotched-tabby pattern in 80% of present-day cats (W841X) occurred in our ancient dataset not earlier than the Medieval period in SWA (3%, minimum number of total alleles, see methods) (Fig. 2). Thereafter its frequency increased in Europe, SWA, and Africa (50% in total) showing late expansion of this typically domestic allele.

**Figure 2.**
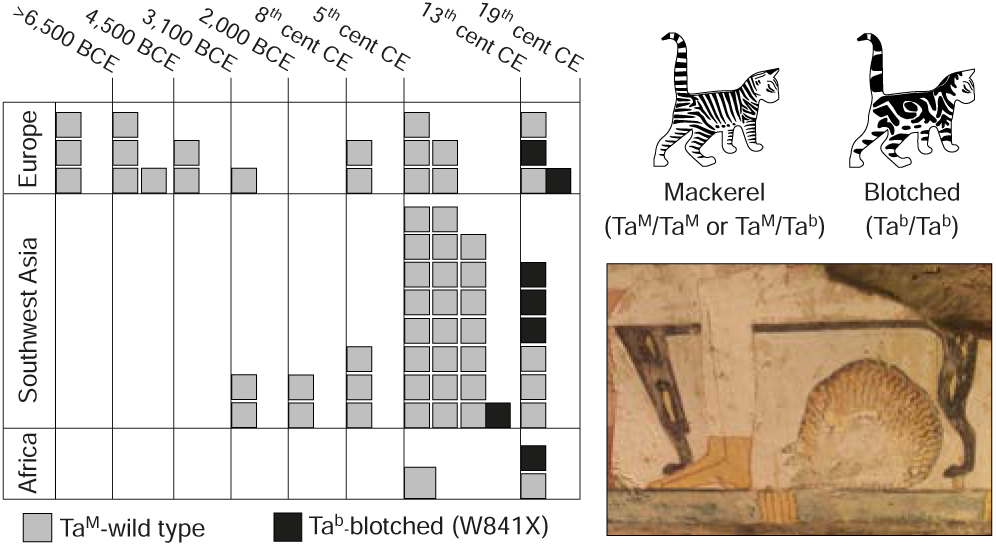
Spatiotemporal representation of the alleles determining the phenotypic variation in the shape of tabby patterns, mackerel and blotched (23). To overcome issues of potential allelic drop-out, each individual is defined by at least one observed allele, except for the few instances in which both alleles were detected. The image below shows a “cat under the chair” with a tabby mackerel marking, typical of *F. s. lybica* (Anna (Nina) Macpherson Davies, *Copy of wall painting from private tomb 52 of Nakht, Thebes (I, 1, 99-102) cat eating fish,* 20th century (1901 - 2000), paint, tempera AN1947.42; Ashmolean museum, Oxford, UK; Image © Ashmolean Museum, University of Oxford).

## Discussion

### Origin and dispersal of domestic cats

Our data show that mitotype IV-A* had a wide distribution stretching across Anatolia from west to east throughout the Neolithic, Bronze Age, and Iron Age. Its range may have extended as far south as the Levant, where we inferred the presence of subclade IV-B. These findings suggest that in the Fertile Crescent, cats developing a commensal relationship with early farming communities during the early Neolithic carried at least the mitotypes IV-A* and IV-B. Mitotype IV-A* later spread to most of the Old World, representing the Near Eastern contribution to the mtDNA pool of present-day domestic cats. This spread may have started as early as ca. 4,400 BCE into Southeast Europe, the date of IV-A*’s first appearance in our European dataset, and thus subsequent to the neolithisation of Europe. This suggests that the human-mediated translocation of cats started already in prehistoric times, corroborating the interpretation of the finding of a cat buried ca. 7,500 BCE in Cyprus (11). We also found IV-A* in cat remains from the Roman-Egyptian port of Berenike on the Red Sea (Fig. 1a-c), which may hint at an introduction of cats from SWA or from the eastern Mediterranean to Egypt.

For the first time we provide evidence for an African contribution, namely clade IV-C, to the mitochondrial gene pool of domestic cats. Indeed, we found the lineages C1 and C* in the majority of Egyptian cat mummies. These cats were worshipped and, during the Greco-Roman period, kept in temple precincts and after their death mummified and deposited in sacred places (18). We show that, despite a local ban on cat trading being imposed in Egypt as early as 1,700 BCE (26), cats carrying IV-C mtDNA spread to most of the Old World. Their increasing popularity among Mediterranean cultures – including Phoenicians, Carthaginians, Greeks, Etruscans, and Romans – and particularly their usefulness on ships infested with rodents and other pests presumably triggered their dispersal across the Mediterranean (26). Indeed, depictions of cats in domestic contexts, already frequent during the New Kingdom in Egypt ca. 1500 BCE (“cat under the chair”, Fig. 2), are found on Greek artifacts from as early as the end of the 6^th^ century BCE (SI Methods). The Egyptian cat must have been very popular, as IV-C1 and C* represented more than half of the maternal lineages in Western Anatolia during the 1^st^ millennium CE, and occurred twice as frequently as the local mitotype IV-A*. This suggests that the Egyptian cat had interesting properties, presumably acquired during the tightening of the human-cat relationship that developed during the Middle and New Kingdoms and became even stronger afterwards (1, 18). Since the most significant genetic changes that distinguish wild and domestic cats are apparently linked to behavior (19), it is tempting to speculate that the success of the Egyptian cat is underlain by changes in its docility, making it more attractive to humans.

North of the Alps, domestic cats appeared soon after the Roman conquest, yet remained absent outside the Roman territory until Late Antiquity (2). In medieval times it was compulsory for seafarers to have cats onboard their ships (27), leading to their dispersal across routes of trade and warfare. This evidence explains, for example, the presence of the Egyptian lineage IV-C1 at the Viking port of Ralswiek (7^th^-11^th^ century CE) (27). The expansion of the domestic cat may have been fostered by a diversification in their cultural usage, which in Medieval Europe included the trade of domestic cat pelts as cloth items (28). Spread of the black rat (*Rattus rattus*) and house mouse (*Mus musculus*) by sea routes as early as the Iron Age, documented by zooarchaeological and genetic data (3), likely also encouraged cat dispersal for the control of these new pests.

Increasing translocation as a result of long-distance trade is also witnessed by the finding of Asian *F. s. ornata* mtDNA in cats from the aforementioned Roman Red Sea port of Berenike (1^st^-2^nd^ century CE) and from Turkey in the 6^th^-7^th^ century CE. This was likely the result of increasingly intensive and direct trade connections between South Asia and the Mediterranean basin via the Indian Ocean and Red Sea (29), but possibly also via the Silk Road connecting Central Asia with Anatolia (30). Long-distance maritime routes (31), as described for instance in the 1^st^ century AD *Periplus of the Erythraean Sea*, likely explain the occurrence of IV-A*, typical of SWA, as far south as East Africa (#30, Fig. 1a-c).

Upon arrival in these various new locations, introduced cats reconfigured the phylogeographic landscape of the species through admixture with local tame or wild cats, leading to a transfer of deeply divergent mitochondrial lineages in the domestic pool (IV-D/E and III *F. s. ornata*). Modern genetic data have shown that admixture with domestic cats still occurs today in European wildcat populations (13), and intensive conservation programs have been implemented to preserve the integrity of *F. s silvestris* (5, 32).

### F. s. lybica in ancient Anatolia and Southeast Europe

Our aDNA data also shed light on the past distribution of wildcats in Eurasia (Fig. S4), with interesting implications for ecology and conservation biology. A clear understanding of the present distribution of wild *F. s. lybica* in Anatolia has proven elusive until now due to a lack of genetic data. It has commonly been assumed that the native range of the modern European wildcat includes Anatolia (6), based on skull and coat features (33) and biogeographic models (34). Our phylogeographic reconstruction demonstrates that mtDNA clade IV, corresponding to *F. s. lybica*, was predominant in Anatolia for many millennia beginning in the Neolithic at the latest. Not a single instance of clade I, corresponding to *F. s. silvestris*, was detected in our samples from SWA (Fig. 1). Nevertheless, we cannot exclude its presence in the wild of Anatolia, in particular in forest and mountain refuges of Northern Anatolia and the Caucasus (up to 22%, 95% Bayesian confidence interval, N=14 in pre-Classical Anatolian sites).

Cats carrying IV-A1 mtDNA were present in Southeast Europe by the beginning of the 8^th^ millennium BCE. This suggests that *F. s. lybica* was distributed across Anatolia from the early Holocene at the latest, prior to the present-day conformation of the Black Sea, and that it made its way to Southeast Europe before the onset of farming in the Neolithic. *F. s. silvestris* and *F. s. lybica* occur across different biotopes that include, respectively, temperate woodland and open bushland (5). The expansion of open bushlands during the Late Pleistocene might have attracted *F. s. lybica* into the Balkans when the Bosporus was a land bridge and the Balkans represented a refuge for warm-adapted species (e.g.; (35, 36). A split in an ancestral Anatolian cat population in the late Pleistocene, presumably during the Last Glacial Maximum, followed by local differentiation and/or drift and founder effect, might have been responsible for the distribution of distinct clade IV mtDNA lineages in Anatolia and Southeast Europe. Currently, IV-A1 is found in the European wildcat population and also in modern domestic cats (Fig. 1,(6)). Our data imply that admixture episodes potentially occurring through time between overlapping populations of wild *F. s. silvestris* and *F. s. lybica* could be in part responsible for *F. s. lybica* introgression in present-day European wildcat populations. Therefore, conservation programs should also take into account past natural admixture when aiming at neutering and removing hybrids that are believed to cause cryptic extirpations of wild *F. s. silvestris* populations (5).

### Evolution of the tabby pattern

Our study also sheds light on the late emergence in domestic cats of a key phenotypic trait, the blotched coat marking caused by a point mutation in the *Taqpep* gene (23). Wildcats exhibit a mackerel-like coat pattern, whereas the blotched pattern is common in many modern domestic breeds (23). In our dataset, the first occurrence of the recessive allele W841X associated with the blotched markings dates to the Ottoman Empire in SWA, and later increases in frequency in Europe, SWA, and Africa (Fig. 2). This result is in agreement with the iconography from the Egyptian New Kingdom through the European Middle Ages, where cats’ coats were mainly depicted as striped, corresponding to the mackerel-tabby pattern of the wild *F. s. lybica* (1, 18) (Fig. 2). It was only in the 18^th^ century CE that the blotched markings were common enough to be associated with the domestic cat by Linnaeus (23), and physical traits started to be selected only in the 19^th^ century CE for the production of fancy breeds (25). Thus, our data suggest that cat domestication in its early stages may have affected mainly behavioral features as indicated by recent genomic data (19). Thus, selective pressure appears to have been low in the past and distinctive physical and aesthetic traits may have been selected only recently. A similar pattern of late emergence of other phenotypic traits has been observed in chicken (37), but contrasts with what has been observed in horses, where coat color differentiation appears at an early stage of domestication (22).

## Conclusions

The aDNA study we present, the first comprehensive genetic study of cats across time and space, provides answers to longstanding questions concerning the domestication process of the cat. By revealing the original phylogeographic distribution of wildcats and its profound modification through human-mediated dispersal of tamed cats through time, we show that both Near Eastern and Egyptian cat lineages contributed at different times to the maternal genetic pool of domestic cats, with each representing the vast majority in present-day cat breeds. Our results indicate that cat domestication was a complex long-term process with extensive translocations that allowed admixture events between geographically separated *F. s. lybica* populations and other cat subspecies (e.g. *F. s. ornata*) at different points in time. The pattern of dispersal of mtDNA haplotypes that emerged from our ancient dataset provides evidence for both human movements along maritime and terrestrial routes of trade and connectivity, and concomitant human-driven dispersal of cats from Egypt by Classical times.

## Methods

### Ancient DNA analyses

Ancient DNA analysis was performed in dedicated aDNA facilities in Paris and Leuven from bone, teeth, skin, and hair samples (the last two when available in Egyptian mummies) of 352 ancient cats. The ages of the archaeological remains were determined using direct accelerator mass spectrometry (AMS) radiocarbon dating (KIK-IRPA, Belgium), stratigraphic associations with AMS dates, and contextual archaeological evidence (Dataset S1). All dates in the text are reported in calibrated radiocarbon years BCE. DNA was also extracted from claws and skin samples of 28 modern wildcats from Bulgaria and East Africa (SI Methods).

Amplification of nine mtDNA and three nuclear DNA fragments in the *Taqpep* gene was preceded by the elimination of carry-over contamination based on the dUTP/UNG system (38) and carried out in three separate multiplex PCRs. Phylogenetically informative SNPs in the mtDNA were selected following the most up-to-date worldwide cat phylogeny (6) (Fig. S1).

We screened the diagnostic SNPs through Pyrosequencing assays (Biotage, Qiagen) and sequencing on a PGM Ion Torrent platform (Institut Jacques Monod, Paris) of amplicon libraries following the aMPlex Torrent workflow and downstream sequence analysis with a bash script described elsewhere (24) (Annexes 1-3, SI). We gathered in total 209 mtDNA profiles ranging from 286 to 449 bp, 12 of which were incomplete profiles generating from two to seven mtDNA fragments. More details about DNA extraction, amplification, sequencing, and the authentication criteria are described in the SI Methods and Tables S1–S4.

### Phylogeographic analyses

Each specimen was assigned to a mtDNA clade using the terminology previously proposed (6), including specimens with an incomplete profile (shapes with an inner dashed line in Fig. 1c). Due to our streamlined sequencing assay, some of the subclades and lineages of IV-A and IV-C observed in the 2007 study were collapsed into a single haplotype, which we named IV-A* and IV-C*, respectively (Fig. S1). The ancient and modern sequences generated here were aligned to 159 sequences from Driscoll’s study (6). A Bayesian tree of 66 unique 286 bp-long haplotypes (Fig. S3) was constructed as described in details in SI. An ML tree, obtained as described in the SI, had the same topology. Frequencies of haplotypes A* and A1 in Anatolia and Southeast Europe, and of clade C in pre- and Roman/post-Roman times in SWA, were tested using a Fisher’s exact test.

### Nuclear markers

We typed allelic variations within the *Taqpep* gene associated with coat color pattern differences − W841X, D228N, and T139N (23). The results presented here are intended to be indicative of allele frequencies. Given the low level of independent replications of our assay and the risk of allelic dropout, especially in ancient degraded samples, we could not ascertain genotypes, except for a few heterozygous samples showing a fairly high number of reads in at least two independent amplifications (Fig. 2 and SI Methods). Assuming that none of the alleles is amplified preferentially, and adopting a conservative strategy accounting for the minimum number of alleles observed, our data across the spatial and temporal framework showed that 7 out of 67 successfully amplified cats possessed at least one mutant Tabby-W841X allele (Fig. 2, and SI Methods), of which two were heterozygotes (BMT2 and MET9). In 88 cats we could screen the allele D228N and in all instances we observed the wild-type. Among 63 cats successfully screened for T139N, we detected the mutant allele (C to A) in three specimens.

## Acknowledgments

This research has been funded by the IAP program (BELSPO), the KU Leuven BOF Centre of Excellence Financing on CAS, and the CNRS (TG & EMG). The sequencing facility of the Institut Jacques Monod, Paris, and JD, were supported by grants to TG from the University Paris Diderot, the “Fondation pour la Recherche Médicale” (DGE20111123014), and the “Région Ile-de-France” (grant 11015901). CO was supported by the FWO mobility program (V4.519.11N, K2.197.14N, K2.057.14N). Faunal research carried out by J. Peters and team at Aşıklı Höyük received funding by the German Research Foundation (DFG PE424/10-1,2). Research by NB, MP, and AC was supported by an ERC grant (#206148) and UK NERC Radiocarbon Facility grant (NF/2012/2/4). Zooarchaeological analyses conducted by A. Bălăşescu at the sites of Hârşova, Vităneşti and Cheia was supported by the Romanian National Authority for Scientific Research, UEFISCDI (PN-II-ID-PCE-2011-3-1015). We thank Greger Larson and E. Andrew Bennett for critical reading of the manuscript; the Ufficio Beni Archeologici della Provincia Autonoma di Bolzano for granting access to the archaeological material of Galgenbühel/Dos de la Forca and Jacopo Crezzini for help in sampling; Marie-Anne Félix, Institut Jacques Monod and École Normale Supérieure, Paris, for granting access to the pyrosequencer; Maarten Larmuseau and Anneleen Van Geystelen for discussions and assistance with nuclear SNP analyses; Katleen Knaepen, Monique Coomans, and Alice Giucca for support in laboratory procedures in Leuven; and Dr. Johan Nackaerts of the veterinary hospital Kruisbos (Wezemaal, Belgium) for providing cat blood samples. We also thank the curators of the following collections for facilitating access to the material under their care and the permission to take tissue samples: the Royal Museum for Central Africa (Tervuren, Belgium), the Muséum national d’Histoire naturelle and Musée du Louvre (Paris), the British Museum and Natural History Museum (London), and the Bavarian State Collection of Anthropology and Palaeoanatomy, Munich.

## Supporting Information

### Precautions taken to minimize contamination

The genetic analyses were performed in the Laboratory of Forensic Genetics and Molecular Archaeology in Leuven (Department of Imaging & Pathology, University of Leuven, Belgium) and in the laboratory of Paleogenomics and Epigenomics of the Institute Jacques Monod (IJM) in Paris. Both laboratories are equipped with dedicated pre-PCR ancient DNA (aDNA) facilities physically separated from post-PCR facilities. The aDNA facility in Paris is under positive air pressure, its equipment and the procedures followed are described in Bennett *et al.,* (2014) (1). In Leuven, access to the pre-PCR laboratory was restricted to a limited number of people and only after wearing clean overalls, gloves, over-shoes, surgical facemasks, plastic spectacles, and following an irreversible sequence of work steps to avoid contamination. Access to the pre-PCR lab in Leuven was not permitted if PCR products had been handled the same day.

The aDNA facilities were routinely cleaned with bleach and RNAse Away (Molecular BioProducts, San Diego, CA, USA) and every item entering the room was extensively washed with bleach or RNAse Away and when possible UV-irradiated.

Buffer, MgCl_2_ and BSA used for preparation of amplification reactions were UV-irradiated to minimize the risk of contamination by animal DNA in reagents (2) and either γ-irradiated water or autoclaved nuclease-free water (Promega) were used. In addition, carry-over prevention strategies based on the use of uracil-N-glycosylase (UNG) were adopted (3).

In Leuven, extractions were performed in a UV-irradiated workstation whereas preparation of amplification reactions was carried out in a UV-irradiated laminar flow cabinet (Esco, Breukelen, Netherlands). In Paris, extractions were done in a UV-irradiated laminar flow hood and amplification reactions were prepared in a dedicated room of the high containment laboratory in a UV-irradiated workstation (Template-Tamer PCR workstation, Qbiogene, Cambridge, U.K.)

In both laboratories, when the two-step nested PCR strategy was adopted (see below), strict temporal and spatial separation of the various steps was adopted to minimize contamination (4). Preparation of second-step nested PCR amplification reactions was carried out in pre-PCR facilities normally used for modern DNA samples, under UV-irradiated workstations or laminar flow cabinets. Multiplex-PCR products of the first amplification step were loaded into the second-step amplification reactions in a laboratory physically separated from pre- and post-PCRs facilities inside a UV-irradiated workstation. Extraction of modern DNA samples was performed in Leuven by a different operator and carried out in pre-PCR laboratories physically separated from the aDNA facilities.

### Preparation of archaeological samples

To extract DNA from teeth and bones, one sample was prepared at a time. Samples were subjected to the following decontamination procedures. The outer surface of bone and teeth samples was removed through sterile blades or, in Leuven, with a Dremel drill (Dremel, Racine, WI, USA).

Additionally, the surfaces of the teeth were gently wiped with 10% bleach and rinsed with bidistilled water. Bone and teeth samples were subsequently ground into a fine powder with a Dremel drill, a mortar or in a 6750 Freezer Mill (SPEX CertiPrep, Metuchen, NJ, USA) and stored at 4°C until DNA extraction. Grinding vials were washed and decontaminated using RNAse Away (Molecular BioProducts, San Diego, CA, USA) and subsequent UV-irradiation (254 nm in cross-liker). In some instances, residues of wrapping tissue were present in skin and hair sampled in Egyptian mummies and they were removed with sterile blades and tweezers.

### DNA extraction and purification

#### Ancient samples

DNA extractions and purifications were performed on aliquots of 100-300 mg of bone or tooth powder, hair tuft (16-102 mg) and skin (up to 250 mg). Bone or tooth powder was incubated for 24-48h at 37°C on a rotating wheel in 1.8 mL digestion buffer containing 0.5 M EDTA (Sigma Aldrich), 0.25 M Na_2_HPO4^3−^ (Sigma Aldrich) and 1% 2-mercaptoethanol (Sigma Aldrich) at pH 8, or in 1.8 mL digestion solution of 0.5 M EDTA pH 8 (Invitrogen, Carlsbad, CA, USA) and 0.25 mg/mL proteinase K (Roche, Penzberg, Germany). Skin and hair tuft samples of Egyptian mummies were incubated for 24h at 56°C on a rotating wheel in 1.8-3.6 mL digestion solution of Tris-HCl 100 mM pH 8, NaCl 100 mM, CaCl_2_ 3 mM, N-Lauroylsarcosine 2%, DTT 40 mM and 0.4 mg/mL proteinase K (5).

After pelleting, DNA was purified following a protocol based on the QIAquick Gel Extraction kit (Qiagen, Hilden, Germany), including additional washing steps with 2 mL QG binding buffer and 2 mL PE wash buffer (Qiagen) (1). Total volumes of 8–36 mL of extract and binding buffer were passed through the silica columns on a vacuum manifold (Qiagen) using 15-25 mL tube extenders (Qiagen). Final DNA elution was done in two steps, each using 27 μL EB elution buffer (Qiagen) heated to 65°C. Each independent extraction batch contained one blank control every five archaeological samples.

All ancient samples were analyzed either by pyrosequencing after nested PCR with biotinylated primers or with Ion Torrent sequencing (see below).

#### Modern samples

DNA extractions and purifications of skin and claw samples from modern wildcats were done in the Laboratory of Forensic Genetics and Molecular Archaeology in Leuven in laboratories that are physically separated from the aDNA facilities and in workflows temporally separated from the aDNA analyses. Samples were directly incubated for 1 hour at 56 °C in 100 μL of incubation buffer (Tissue and Hair Extraction kit, Promega), 0.1 M DTT (Invitrogen) and 2 mg/mL proteinase K (Roche, Penzberg, Germany). After removing the claw or the piece of skin, 300 μL of lysis buffer (DNA IQ Casework Pro Kit for Maxwell 16, Promega) and 3 μL of DTT 1M were added to the solution and purified in a Maxwell 16 (Promega) with a DNA IQ Casework Pro Kit.

All samples were analyzed in Leuven by pyrosequencing after nested PCR amplification with biotinylated primers (see below).

### Selection of SNPs and primer design

#### Mitochondrial DNA

For primer design, we relied on the most up-to-date worldwide cat phylogeny described by the 2007 study of Driscoll *et al*. (6), in which 2604 bp of the ND5, ND6 and CytB genes of the mtDNA were sequenced revealing five different clades (I to V) that correspond to five *Felis silvestris* subspecies (Fig. S1a). The haplotypes from Driscoll’s study were downloaded from GenBank, aligned in Geneious 6 (http://www.geneious.com (7)), against a reference sequence (8), and a maximum-likelihood tree was constructed using PhyML (9) in Geneious. With the aim to design a molecular assay that was suitable for the analysis of ancient degraded samples and at the same time preserved a fairly good phylogenetic resolution at the level of the major clades and subclades, we selected 42 informative SNPs distributed across nine short regions of the ND5, ND6 and CytB genes of the mtDNA. Primer design and multiplex testing were performed following the strategy of the aMPlex Torrent workflow (10). We used the software Primer3 within Geneious to design primers which hybridize to conserved sequences flanking short phylogenetically informative sequences containing one or several diagnostic SNPs. For primer optimization, we also used the software Oligo 6 (Molecular Biology Insights, Inc) to select primers with a temperature of dissociation (Td) of 65 to 67°C (nearest neighbor method) and for which there was no predicted 3’-dimer having a ΔG below −1.6 kcal/mol. In this way, we designed various candidate primer pairs targeting nine regions (<130 bp) containing the diagnostic SNPs selected and tested them for efficiency and dimer formation using quantitative real-time PCR (qPCR) with a LightCycler 480 (Roche). qPCR amplification was carried out with 2 µL of cat DNA (extracted from hairs of a modern cat) at five different concentrations (serial dilutions from 12,5 ng to 20 pg), or 2 µL of water for non-template controls (6 replications per primer pair), in a final volume of 7 µL containing SYBR Green I Master Mix 1x and 0.5 µM of each primer in a 384-well plate with an EpMotion 5070 robot (Eppendorf). The cycling program used was as follows: 5 minutes of polymerase activation at 95°C, followed by 60 cycles at 95°C for 10 seconds and 60°C for 40 seconds. A melting curve was established at the end of the run through a melting step (95°C for 5 seconds, then 65°C for 1 minute and a temperature increase of 0.11°C with continuous fluorescence measurement). We selected primer pairs whose efficiency was above 90 % and that yielded no primer-dimers before 40 cycles. The final primers selected amplified fragments of 60 to 111 bp in length (see Table S1).

In order to choose the best combination of primers to use in multiplex PCRs, we predicted *in silico* their tendency to form primer-dimers using the PriDimerCheck software (http://biocompute.bmi.ac.cn/MPprimer/primer_dimer.html), (11). Based on the output of the program we divided the nine primer pairs in two different multiplex reactions to minimize the predicted stability of 3’-dimers. Each multiplex was optimized by testing different MgCl_2_ concentrations (2 mM, 3 mM and 4 mM) and was carried out with 5 µL of 10 pg/µL cat DNA in a 50 µL reaction volume with a final composition of 1X FastStart High Fidelity Reaction Buffer and MgCl_2_ 3 mM (Roche), 0.25 mM each of dGTP, dATP, dCTP, 0.5 mM of dUTP (Bioline), BSA 1 mg/mL, 0.3 units of Uracil-N-glycosylase (UNG, ArcticZymes), 1.7 units of FastStart Taq polymerase (Roche), 0.15 µM of each primer (Sigma, IDT) and, when appropriate, additional MgCl_2_ to achieve a final concentration of either 3 or 4 mM. The MgCl_2_ solution, reaction buffer and BSA were decontaminated by exposing the solutions in UV-pervious tubes (Qubit^®^, Life Technologies) to UV light for 10 minutes at a short distance as described elsewhere (2). Negative controls for each multiplex reaction were processed in the same way as the samples throughout the whole experimental procedure. The cycling program consisted of 15 minutes at 37°C (carry-over contamination prevention through digestion by UNG of dUTP-containing amplicons), 95°C for 10 minutes (inactivation of UNG and activation of the Fast Start DNA polymerase), followed by 35 cycles at 95°C for 10 seconds and 60°C for 1 minute and a final extension step at 72°C for 10 minutes.

After the multiplex PCR amplification, qPCR analyses of the amplification products were performed using each primer pair individually, to measure the production of each PCR product and of possible primer-dimers involving the specific tested primer pair. Each simplex qPCR was carried out as described above using 2 µL of the multiplex product diluted to 1/50th. In order to minimize carry-over contamination, the dilution of the PCR product was performed in a dedicated laboratory, physically separated from the high containment laboratory, the modern DNA facility, the post-PCR facility and the genomic platform. In this way, we established that a MgCl_2_ concentration of 3 mM was optimal for the multiplex amplification. The sensitivity of the two multiplex PCRs was finally tested by amplifying 20 pg of cat DNA, showing amplification of all fragments, and 5 pg of DNA, in which amplification of some fragments was not obtained.

To assess the phylogenetic consistency of our molecular assay, we constructed a Bayesian tree from the haplotype data of Driscoll et al. (6) reduced to the minimum sequence length generated by our assays (i.e. 286 bp through pyrosequencing, see below) using Mr Bayes (12). The tree preserved the same topology as that previously described in Driscoll *et al*.’s study in all its major clades and subclades with fairly high statistical support in the main nodes (see Fig. S1). The same results were observed with the Maximum-Likelihood method as implemented in PHYML (9) adopting various statistical supports for node definition.

#### Nuclear SNPs in the Taqpep gene

During felid skin development, the loss of function of the *Taqpep* gene, which encodes the protein Tabulin, causes a loss of coat color pattern periodicity, determining the phenotypic variation in the shape of tabby patterns – mackerel (Ta^M^/Ta^M^, Ta^M^/Ta^b^) and blotched (Ta^b^/Ta^b^). Blotched cats were found to carry a nonsense mutation (W841X, G-A transition) in exon 17 of the *Taqpep* gene (13). A second less frequent Ta^b^ allele (D228N, G-A transition) was found to cosegregate with the blotched phenotype in a research colony, whereas a variant at codon 139 (T139N, C-A transversion) was associated to an atypical swirled coat pattern, a phenotype that is similar to the spotting pattern of *Felis nigripes* (13).

The three nuclear SNPs associated to a phenotypic variation in the shape of tabby patters – mackerel (Ta^M^/Ta^M^, Ta^M^/Ta^b^) and blotched (Ta^b^/Ta^b^) (13) – in the nuclear gene *Transmembrane Aminopeptidase Q* (*Taqpep*) were analyzed by amplifying in one multiplex three fragments ranging 64-110 bp in length (see Table S1). Primer pairs were designed and tested as described above for the mtDNA.

### Amplification of DNA

Amplification of the nine mtDNA fragments was preceded by the elimination of carry-over contamination based on the dUTP/UNG system (3) and performed in two separate mixes in a final volume of 30-50 μL, at conditions tested and optimized as described above ‒ 1X FastStart High Fidelity ^Reaction Buffer and MgCl_2_ 3 mM (Roche), 0.25 mM each of dGTP, dATP, dCTP, 0.5 mM of dUTP^ (Bioline), BSA 1 mg/mL, 0.3 units of Uracil-DNA glycosylase (UNG, ArcticZymes), 1.7 units of FastStart Taq polymerase (Roche), 0.15 µM of each primer (Sigma, IDT) ‒ using 3-6 μL of aDNA extract. The following cycle conditions were used: 37°C for 15 min, 95°C for 10 min, 35 cycles of 95°C for 30 sec, 60°C for 1 min, and a final step of 72°C for 4 min.

In a first phase of the project, positive amplification products were reamplified in nested SYBR Green qPCR and submitted to pyrosequencing assays (see below).

In a second phase of the project, amplicon libraries were constructed and sequenced on the PGM Ion Torrent platform of the IJM following the aMPlex Torrent workflow (10) (see below). In order to obtain an approximately even distribution of final reads we tested various primer concentrations for each multiplex. These tests were done with 3.5-350 pg of cat DNA (from FTA blood samples) and were carried out with the same strategy described above by measuring the production of each PCR product through QPCR analyses with each primer pair used individually. The final optimized primer concentrations are reported in Table S1.

The amplification of the three nuclear fragments of the *Taqpep* gene, was done in one separate multiplex PCR with the same conditions as the multiplexes used for the mtDNA amplification except for a higher concentration of MgCl_2_ (4 mM), a denaturation time of 30 sec in the amplification step was used, and 40 amplification cycles. All nuclear amplicons were sequenced exclusively with the PGM Ion Torrent platform.

When necessary, simplex PCR reactions of missing or poorly represented mtDNA fragments were performed using the same PCR conditions as in the multiplex PCR and were sequenced either with pyrosequencing or in the PGM Ion Torrent.

### Pyrosequencing of PCR products

A first analytical phase of the project relied on pyrosequencing assays of 5’-biotinylated PCR products to screen 42 informative SNPs within the mtDNA fragments targeted. Nested PCR primers, sequencing primers and nucleotide dispensation orders (Table S1, S2, S3) were automatically generated with the Assay Design Software v1.0 (Biotage). Pyrosequencing was performed after nested simplex amplification of the nine mtDNA amplicons obtained from two multiplex PCRs. Amplification products were diluted up to 200 times and 2 μL of diluted PCR product were reamplified in nested SYBR Green QPCR simplex reactions using separately ten primer pairs with one 5’-biotinylated primer ‒ one multiplex PCR fragment was covered by two different overlapping primer pairs ‒ in a final volume of 25 µL containing either 1X GoTaq Green Master Mix (Promega) or 1X SYBR Green I Master Mix (Roche) and 0.76 µM of each primer (Table S1). Amplifications were performed in a Lightcycler 480 (Roche) in Paris or in a 7500 Real Time PCR System (Applied Biosystems) in Leuven. The following cycle conditions were used: 95°C for 2 min, 30 cycles of 95°C for 15 sec, 60°C for 1 min, and a final step of 72°C for 4 min. Dimers were identified by subsequent melting curve analysis.

Positive PCR products from the SYBR Green QPCR simplex reactions were processed to yield single-stranded DNA fragments before the pyrosequencing reaction by attachment to streptavidin-coated beads, followed by strand separation and washing steps performed in a Vacuum Prep Workstation following manufacturer instructions (Biotage PyroMark Q96 System and PyroMark Q24, Qiagen). The beads with single-stranded DNA fragments were then transferred to a buffer containing the sequencing primer (Sigma, IDT), which is annealed by heating to 80°C for 2 min and cooling for at least 5 min. Pyrosequencing was performed following the manufacturer’s instructions in a Biotage PyroMark Q96 at the Institut de Biologie de l’Ecole Normale Supérieure in Paris and in a PyroMark Q24 (Qiagen) at the Laboratory of Forensic Genetics and Molecular Archaeology in Leuven. Results and sequences were analyzed in the PyroMark Q24 software (Qiagen) or the PyroMark Q96 software (Biotage). In three instances (museum specimens 2129, 2130 from Kenya and 17153 from Rwanda), out of phase shifts in the pyrograms of one fragment (i.e. fragment 3) were observed suggesting the presence of an additional variable site. To account for the additional SNP (G/A at np 13537), the dispensation order was changed and in all instances the correct phase was recovered.

### Library preparation and Ion Torrent sequencing of amplified products in the Genomic Platform of the IJM

In a second analytical phase of the project amplicon, libraries were prepared from the PCR products of the three multiplex PCR reactions of the ancient samples (nine fragments in two multiplexes of the mtDNA and three fragments in one multiplex of the nuclear *Taqpep* gene) and sequenced in a PGM Ion Torrent platform using the aMPlex Torrent workflow (10). Amplifications, library preparations and Ion Torrent sequencing were all performed at the IJM in Paris.

Amplification products from the three multiplexes were pooled for each individual and blank control in a 96-well plate. We constructed up to nine independent libraries. Each time, samples were amplified in batches of 44 duplicates from one extract, plus eight extraction and PCR controls, using all 96 Ion Torrent barcodes. In some instances, samples were amplified in triplicate or repeated in duplicate in separate amplification batches.

The following steps were performed using a Tecan Freedom Evo 100 robot equipped with a 4 channel liquid handling arm using disposable tips, a gripper to move objects, a double thermoblock and an automated solution for vacuum solid phase extraction.

We first performed end-repair with the NEBNext End Repair module (New England Biolabs) using 40 µL of pooled multiplex PCR (containing no more than 5 picomoles of amplicon products) in a 50 µL reaction volume with 0.1 µL of End Repair Enzymes. The reaction was incubated for 30 minutes at 25°C and then purified using the NucleoSpin 96 PCR clean-up kit as recommended by the manufacturer (Macherey-Nagel ref. 740658). DNA was eluted in 50 µL and 20 µL. Sample-specific Ion Torrent barcoded adaptors (1 µL of annealed A+P1, 20 µM) were ligated using the NEB Next ligation module (New England Biolabs) in a 30 µL reaction volume with 1 µL of Quick Ligase, incubated for 30 minutes at 16°C. After the ligation, 60 µL of NT binding buffer (Macherey-Nagel) was added and all samples were pooled in a tube before purification on a silica column (Qiagen or Macherey-Nagel). Ten µL (out of 30 µL) of the pooled barcoded PCR products were then size selected using the Caliper Labchip XT (Perkin Elmer) by setting a size range of 144-197 bp in order to recover all the amplicons and remove adapter-dimers. Final recovery was done in 20 µL of collection buffer. Size-selected products were subjected to nick-repair and amplification in a 24 µL final volume reaction using NEB OneTaq Hot Start and containing Ion Torrent primers A and P1 (0.5 µM). The reaction was incubated for 20 min at 68°C (nick-repair step), then amplified using the following program: initial denaturation at 94°C for 5 min, (94°C for 15 sec, 60°C for 15 sec, 68°C for 40 sec) for 6 cycles, final elongation at 68°C for 5 min. Products were then purified with a Qiaquick PCR purification kit (Qiagen). The size distribution and concentration of the library was assessed on the Agilent 2100 Bioanalyzer. Emulsion PCR and Ion Sphere Particle enrichment were conducted with the Ion OneTouch System (Life Technologies) using the Ion OneTouch 200 Template kit v2 DL according to the manufacturer’s protocol. Each of the nine DNA libraries was sequenced independently on the Ion Torrent Personal Genome Machine (PGM) Sequencer using the Ion PGM 200 Sequencing Kit and Ion 314 semiconductor sequencing chips (Life Technologies).

### Analysis of sequencing data

Using the software provided in the Torrent suite on the Ion Torrent server, sequencing reads were demultiplexed and mapped with the Torrent Mapping Alignment Program (TMAP) to a reference fasta sequence composed of the 12 fragments (9 mitochondrial, 3 nuclear) separated by spacers of 100 Ns. The quality of the sequencing reaction and of the library analyzed was assessed within the analysis report of the Torrent Browser. Details about the quality of each Ion Torrent run are reported in Table S4.

Downstream sequence analysis was performed by first renaming the BAM and BAI files to allocate simple descriptive names using bash commands. The files were then analyzed with a bash script previously developed (10) (Annex 1, SI) dependent on the following programs (and tested with the versions indicated between brackets): featureCounts (Subread package 1.4.6) (14), SAMtools and bcftools (1.2) (15). BAM files were converted to SAM files with SAMtools. The number of reads per amplicon for each sample was estimated with featureCounts, which uses a GFF file (Annex 3, SI) describing the features of the fasta file (Annex 2, SI) used for mapping. In the gff file we defined the sequences of the PCR products and the primers as “miscellaneous features” to count the proper PCR products as well as the reads that contain either the forward or the reverse primers. This ensures the accurate estimation of the mapped reads while measuring potential primer dimers identified as supernumerary primer-only counts. Finally, SAMtools *mpileup* and *bcftools* were used to generate a consensus sequence of the 12 concatenated fragments obtained for each individually barcoded sample. We used Geneious (http://www.geneious.com, (7)) to import both the consensus fasta and the initial BAM files. Consensus sequences were aligned against a reference sequence (NC001700), with manual editing when necessary. The various outputs were verified by analyzing the initial sequencing reads within the BAM files displayed within Geneious.

By applying the authentication criteria described below, our sequencing strategy made it possible to gather in total 209 mtDNA profiles out of the 352 ancient cats analyzed (Datasets S1, S2). We observed that success rates decreased with increasing environmental temperature. Indeed, only 12 out of 68 ancient Egyptian samples analyzed (Predynastic to Greco-Roman, including 52 mummies) and none of the 14 Neolithic and Bronze Age samples from Cyprus, Jordan and Syria yielded DNA sequences.

In 197 specimens complete mtDNA profiles (i.e. the nine fragments analyzed) were obtained, with total length of sequences read ranging from 286-299 bp in samples sequenced exclusively by pyrosequencing respectively with the Pyromark Q96 (N=13) and Pyromark Q24 (N=23), up to 449 bp in samples entirely sequenced in the Ion Torrent platform (N=150), whereas the remaining samples were sequenced by a combination of both techniques (N=11) (Dataset S2). In 12 samples incomplete profiles were generated by successfully amplifying two to seven mtDNA fragments. In most instances (11/12 samples) the longest fragments ranging between 101-111 bp failed to amplify, as expected for ancient samples following diagenetic trajectories of biomolecular decay.

A tree of 66 unique 286 bp-long haplotypes (Fig. S2) was constructed with MrBayes (12) implementing the HKY85+G model, as identified by ModelTest (16), from two runs of five million generations and four heated chains each. The posterior probabilities and consensus tree of 66 unique 286 bp-long haplotypes (Fig. S2) constructed with MrBayes were visualized with FigTree v1.4 (Fig. S2) (http://tree.bio.ed.ac.uk/software/figtree/). The same topology of the tree was obtained with PhyML (9) with various branch support methods – bootstrap (1000 iterations) and SH.

One of three nuclear fragments could be amplified in 97 samples in at least one amplification reaction. The Tabby-T139A allele could be amplified and sequenced in 63 samples, the Tabby-D228N in 88 samples and the Tabby-W841X allele in 67 samples. The three fragments together were successfully amplified and sequenced in 48 samples.

### Authentication of aDNA data

- In the nine Ion Torrent sequencing runs carried out, 50 blank extractions and PCR controls were processed in parallel with the samples, providing an average coverage of 0.54, ranging from 0.09 to 1.52, for the mtDNA amplicons (with a maximum of five reads observed), and 0.09 ranging from 0 to 0.33 for the nuclear DNA amplicons (with a maximum of two reads). Only in one instance, an extraction control provided from 7 to 20 reads in six mtDNA fragments, a result that nevertheless was not replicated in a second control from the same extraction batch. All successful samples from that sequencing run showed a larger number of reads (>150) and the sequences were replicated in independent experiments. Furthermore, all the successful samples from that extraction batch provided results replicated at high average coverage (>150 reads) from multiple amplification and/or extractions or showed a different sequence from that in the blank control. In the nested PCR approach no extraction or PCR control was ever amplified.

- In all instances, mtDNA haplotypes obtained by Ion Torrent sequencing or pyrosequencing were reproduced in multiple experiments, from at least two and up to eight independent multiplex PCR experiments from one or up to three DNA extracts (e.g. samples BM06 and OXY3). When possible, independent DNA extractions were performed from anatomically distant bone samples or from different tissues (e.g. skin, hair and bone for the mummified samples).

Given the lower chance of successful amplification of nuclear fragments and the extremely low number of reads observed in the negative controls (average read number 0.09), in some instances results could not be replicated in independent amplification and yet considered authentic when at least 20 reads were obtained (see Dataset S2). For this reason and given the degraded nature of aDNA, we cannot exclude that, due to allelic drop-out, our strategy for nuclear SNPs analysis might have underestimated the number of alleles observed for the three SNPs examined. Therefore, we adopted a conservative strategy accounting only for the minimum number of alleles observed.

- Samples in which one or more mtDNA fragments failed to amplify or were poorly represented after multiple attempts of multiplex PCR amplification were re-analyzed using either *i)* simplex PCRs to increase the chances of successful amplification of the missing or poorly represented fragments and sequenced in the Ion Torrent, or *ii)* independent simplex PCRs, followed by nested qPCR with a 5’- biotinylated primer and pyrosequencing.

Samples in which all fragments failed to amplify or that were covered by <10 reads and those in which inconsistencies of the sequences were found across multiple replicates were discarded and considered unsuccessful.

- Of the 150 successful samples with a complete 449 bp-long profile sequenced with the Ion Torrent, in 33 the average coverage obtained from independent replicates of the multiple PCRs was >1000 reads, in 28 samples it was between 500 and 1000, in 71 samples between 50 and 500, and in 18 samples it was between 14 and 50. In the latter case, to ensure higher reliability of poorly covered fragments, simplex PCRs were performed that allowed obtaining up to thousands reads (Dataset S2).

In samples where both Ion Torrent and pyrosequencing techniques were applied on the same fragment the results were always consistent.

- Nuclear fragments were represented in all samples by >10 reads in two or more independent PCR amplifications, or by >20 reads in samples where a single amplification was obtained (Dataset S2). In 73 samples the average coverage of the successful nuclear fragments amplified was >100 reads.

### Iconographic evidence in Europe

Representations of cats in domestic contexts are known from various artifacts and funerary monuments in the Greek world during the 6^th^ c. BC. Here we report a list of some of the iconographic depictions.

- The Arkesilas Cup found in the Etruscan site of Vulci and dated to about 565/560 BC (Cabinet des médailles of the Bibliothèque nationale de France in Paris (inv. 189);
- The funerary statue of Themistocle 510 BC (National Archaeological Museum, Athens); (see also Luce, J-M. (2015) Les chats dans l’Antiquité grecque. In “Chiens & Chats dans la Préhistoire et l’Antiquité”, Bellier, C., Cattelain, L. et Cattelain, P. (eds), Editions du Cedarc, Bruxelles.)
- The red-figure vases of the “Cat and Dog Painter” (The Beazley Archive Pottery Database, BAPD, http://www.beazley.ox.ac.uk/XDB/ASP/default.asp)
- Greek Coins (Kraay, Colin M (1976), Archaic and Classical Greek Coins, New York: Sanford J. Durst); for images see: Lauwers, C. “Des monnaies entre chien et loup” In “Chiens & Chats dans la Préhistoire et l’Antiquité”, Bellier, C., Cattelain, L. et Cattelain, P. (eds), Editions du Cedarc, Bruxelles)

### Datasets and supplementary tables

**Dataset S1.** List of all the ancient samples analyzed in this study. Unsuccessful and successful samples with a complete and incomplete mtDNA profile are indicated in red, green and yellow respectively.

**Dataset S2.** List of samples successfully analyzed in this study and detailed information about the samples, the dating, the genotyping procedures followed, the mtDNA haplotypes and the polymorphic states of the three nuclear markers investigated. For samples analyzed with the aMPlex Torrent approach, the average number of reads obtained in all fragments amplified for the mtDNA and the nuDNA fragments is reported (in samples in which simplex amplifications were performed, the maximum and minimum number of reads obtained from different fragments and/or PCRs reactions is indicated in brackets). The color code associated to sample ID in column B is identical to that used in Dataset1 whereas that used for the clade/subclade in column J is identical to that used in the various trees depicted in Fig.1, S1 and S2.

**Table S1.**
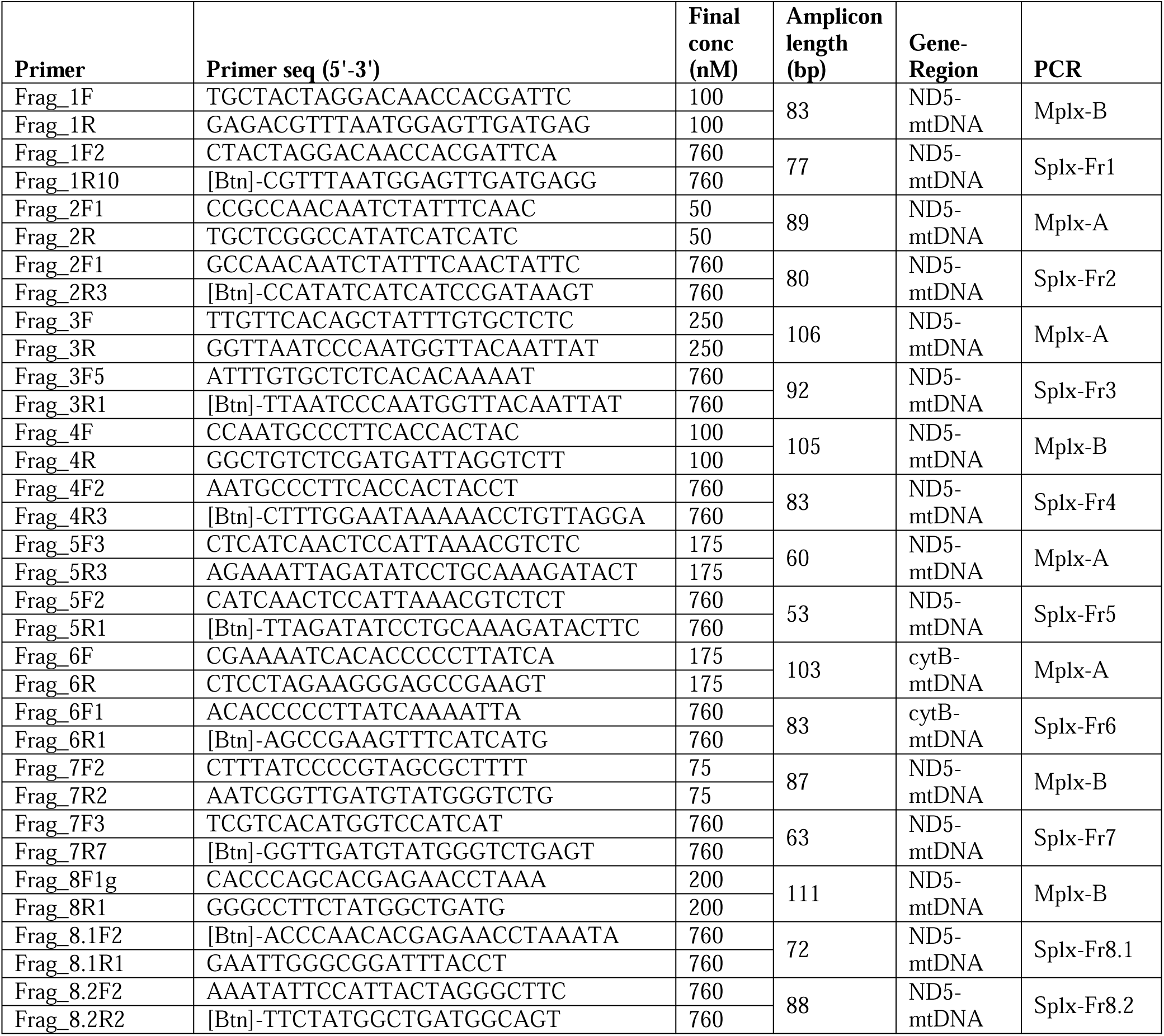
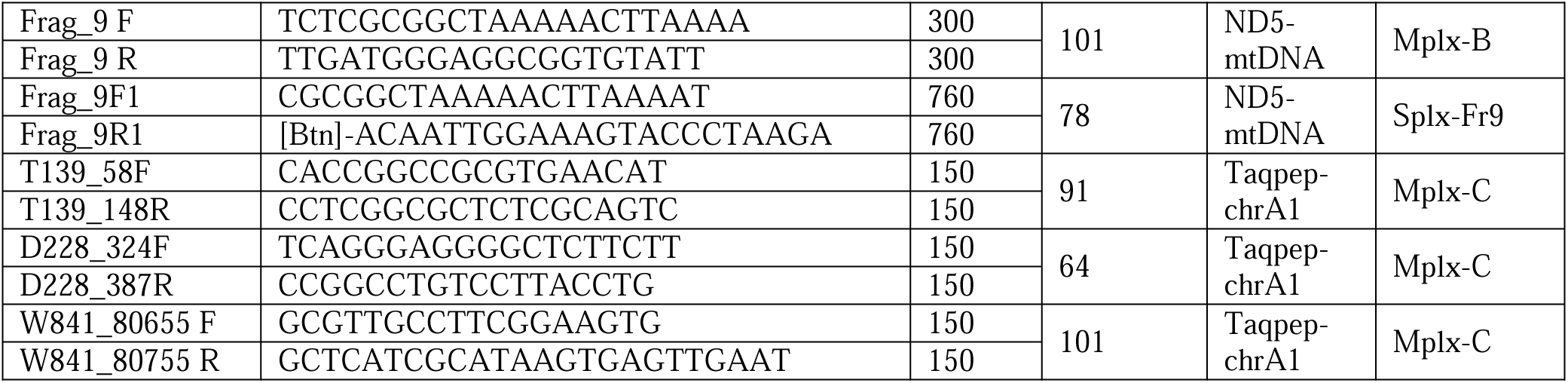
All PCR primers used in multiplex and simplex PCRs in this study

**Table S2.**
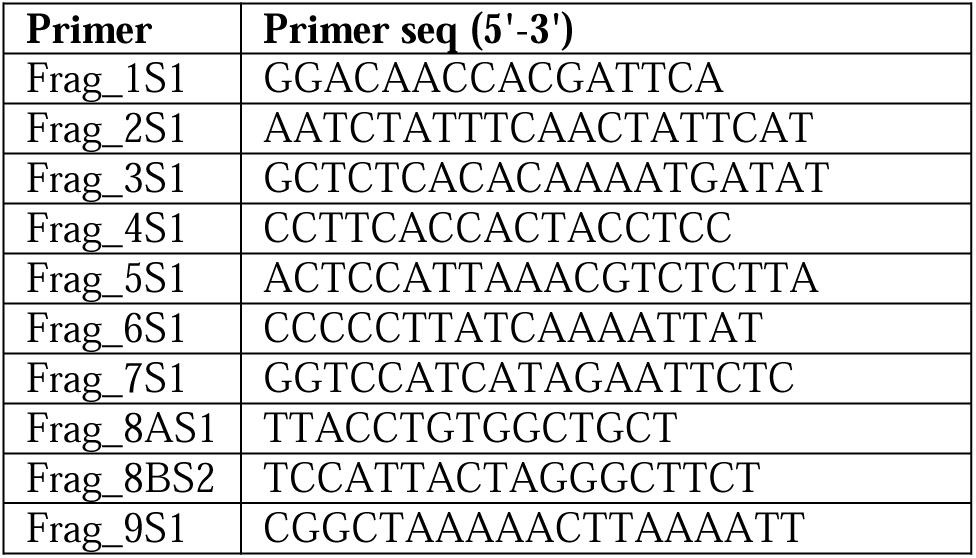
Pyrosequencing primers (mtDNA)

**Table S3.**
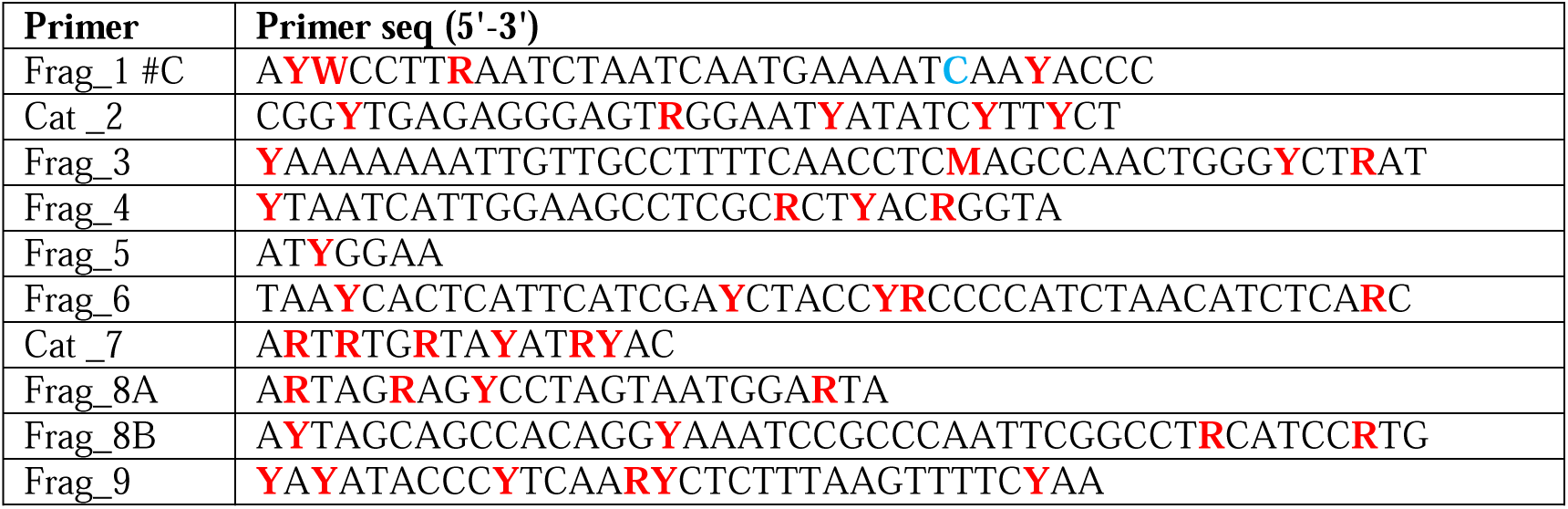
Sequences to analyse in pyrosequencing reactions (mtDNA)

**Table S4.** Details of Ion Torrent sequencing runs

| ION | Barcoded ancient samples (N) | Polyclonal | Total reads | Mean read length (bp) | Aligned reads |
| --- | --- | --- | --- | --- | --- |
| 31 | 88 | 43% | 265,745 | 78 | 136,382 |
| 32 | 88 | 41% | 377,657 | 56 | 28,041 |
| 33 | 88 | 34% | 261,117 | 72 | 148,205 |
| 34 | 88 | 39% | 254,715 | 58 | 24,252 |
| 40 | 80 | 44% | 515,694 | 79 | 343,414 |
| 41 | 88 | 44% | 536,969 | 85 | 414,676 |
| 42 | 88 | 40% | 566,386 | 92 | 325,774 |
| 44 | 88 | 31% | 658,721 | 92 | 494,968 |
| 46 | 80 | 32% | 629,746 | 86 | 376,467 |

**Figure S1.**
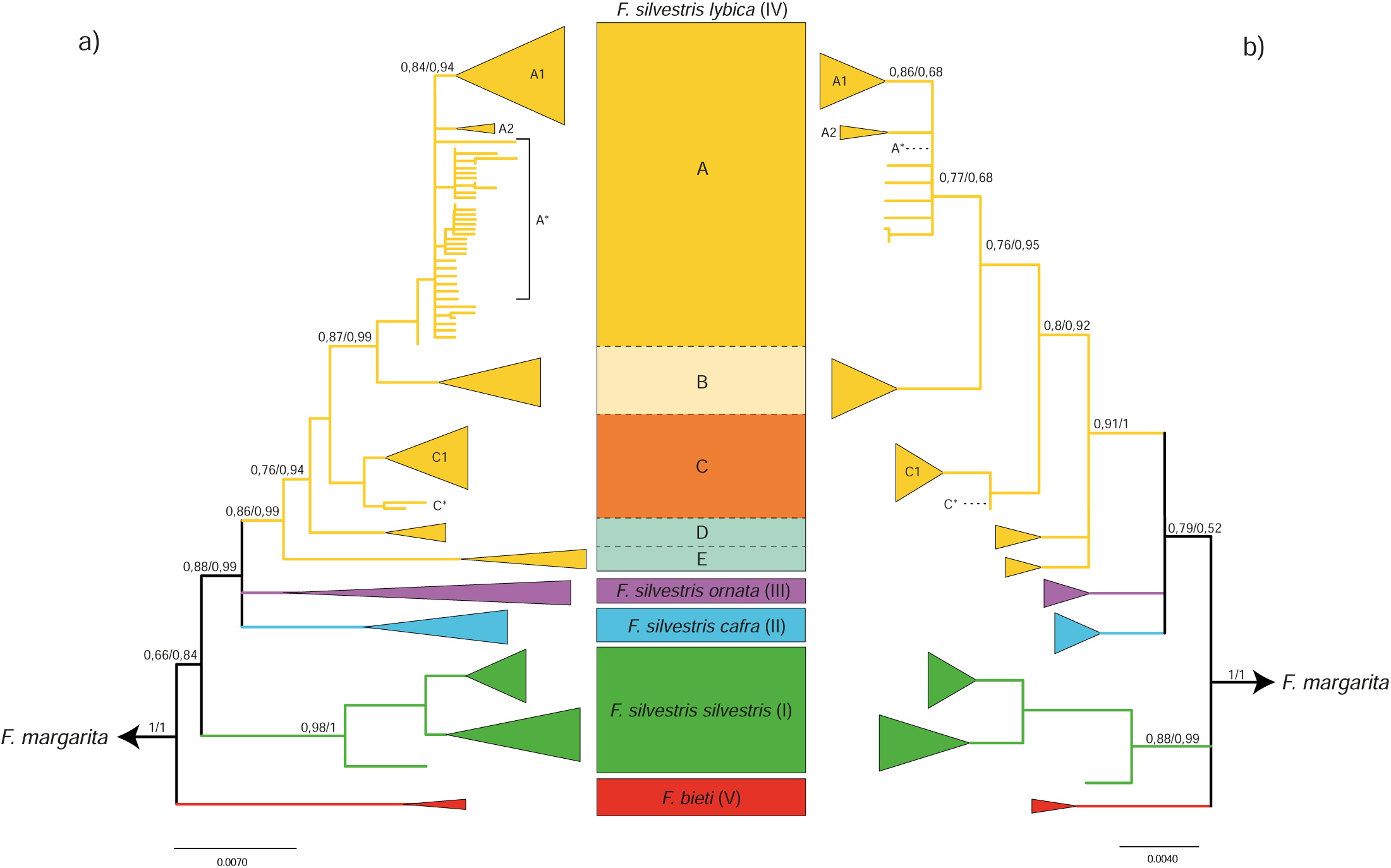
a) Maximum-likelihood (ML) trees of haplotypes from Driscoll *et* al. (6) and b) of the same haplotype data reduced to the minimum sequence length − 286 bp − generated by our assay. Subspecies and clade names as described by Driscoll et al. are reported in the rectangular shape between the trees, and colors of clades and subclades are as in haplotypes of figure 1b. Names of subclades within clade IV (*F. s. lybica*) dubbed in this study are reported in each tree. Some of the subclades and lineages observed in the 2007 study of Driscoll *et al*. were collapsed in a single root haplotype (IV-A*, IV-C*). The ML trees were generated using PHYML3.0 (9, 18) and the Shimodaira-Hasegawa-like branch test (SH) was used to evaluate the statistical support of the nodes. A Bayesian tree generated using Mr Bayes had the same topology and was used to evaluate the posterior probabilities of the nodes. Statistical support for the main nodes is reported as SH/posterior probability values.

**Figure S2.**
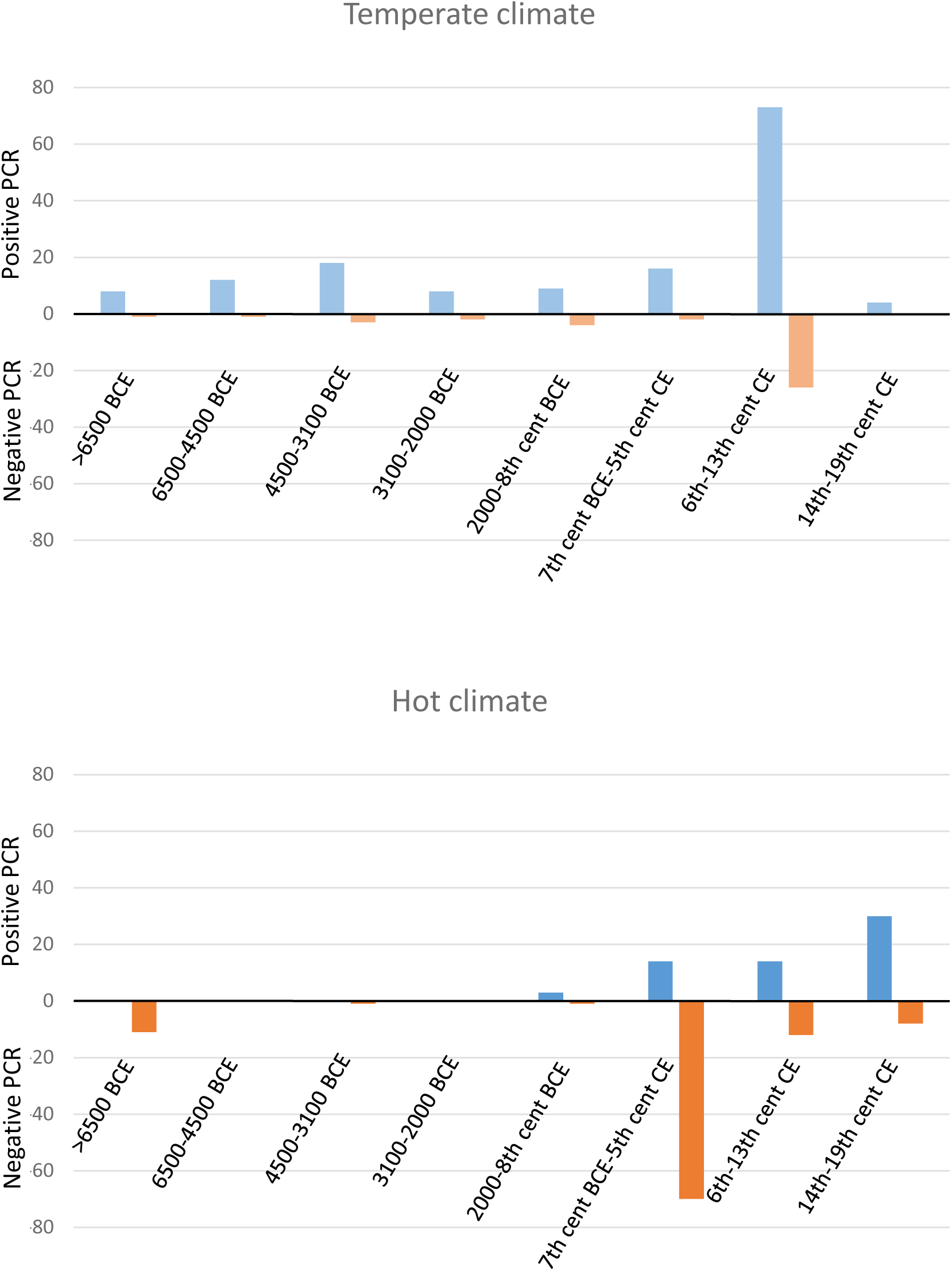
Distribution of the samples allowing or not PCR amplifications according to age and environmental temperature. Samples are classified in time bins as indicated and countries of origin are classified according to climate either as “temperate” on the top graph (Europe, Armenia, Iran and Turkey) or “hot” on the bottom graph (Africa, Jordan, Lebanon, Oman, Saudi Arabia, Syria, Egypt). The number of samples that yielded PCR products in each category are represented in blue above the zero line with a scale oriented upward, whereas the number of PCR negative samples are represented in orange below the zero line with a scale oriented downward.

**Figure S3.**
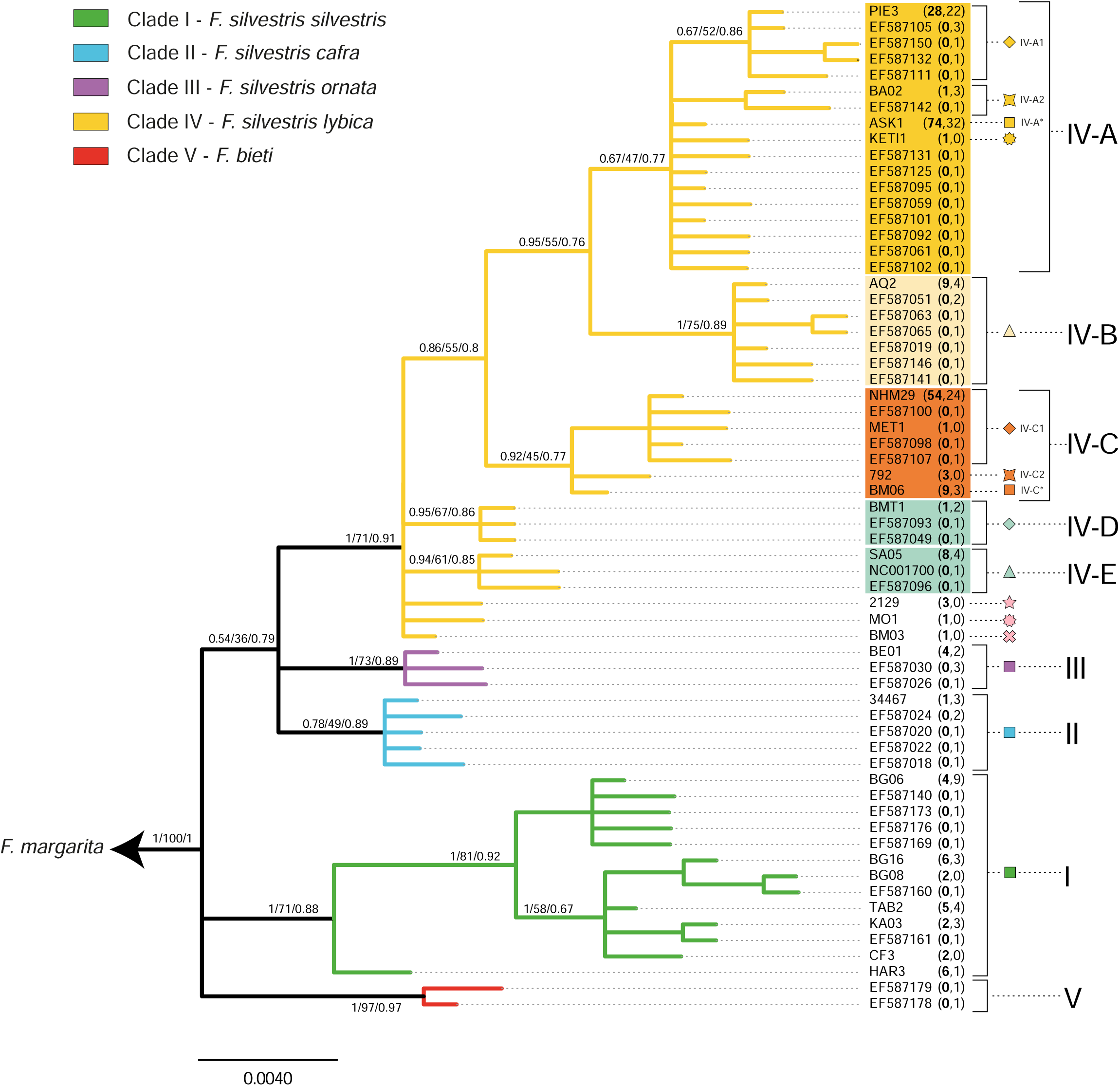
Bayesian tree of 286 bp-long haplotype data from this study and literature (6), represented in a schematic way in Figure 1b. Colors of branches in the five *Felis silvestris* clades are as in figure 1. Statistical support values are reported as Bayesian posterior probability/bootstrap/SH-values obtained by three different methods (Mr Bayes, PHYML3.0, PHYML3.0, respectively) reproducing the same topology. Each haplotype is indicated by the name of one specimen (or a Genbank accession number for sequences from the literature not found in the dataset generated in this study). For each haplotype, the number of sequences from this study (in bold) and from the 2007 study of Driscoll *et al*. (haplotypes deposited in GenBank) sharing the same 286 bp-haplotype is indicated in brackets.

**Figure S4.**
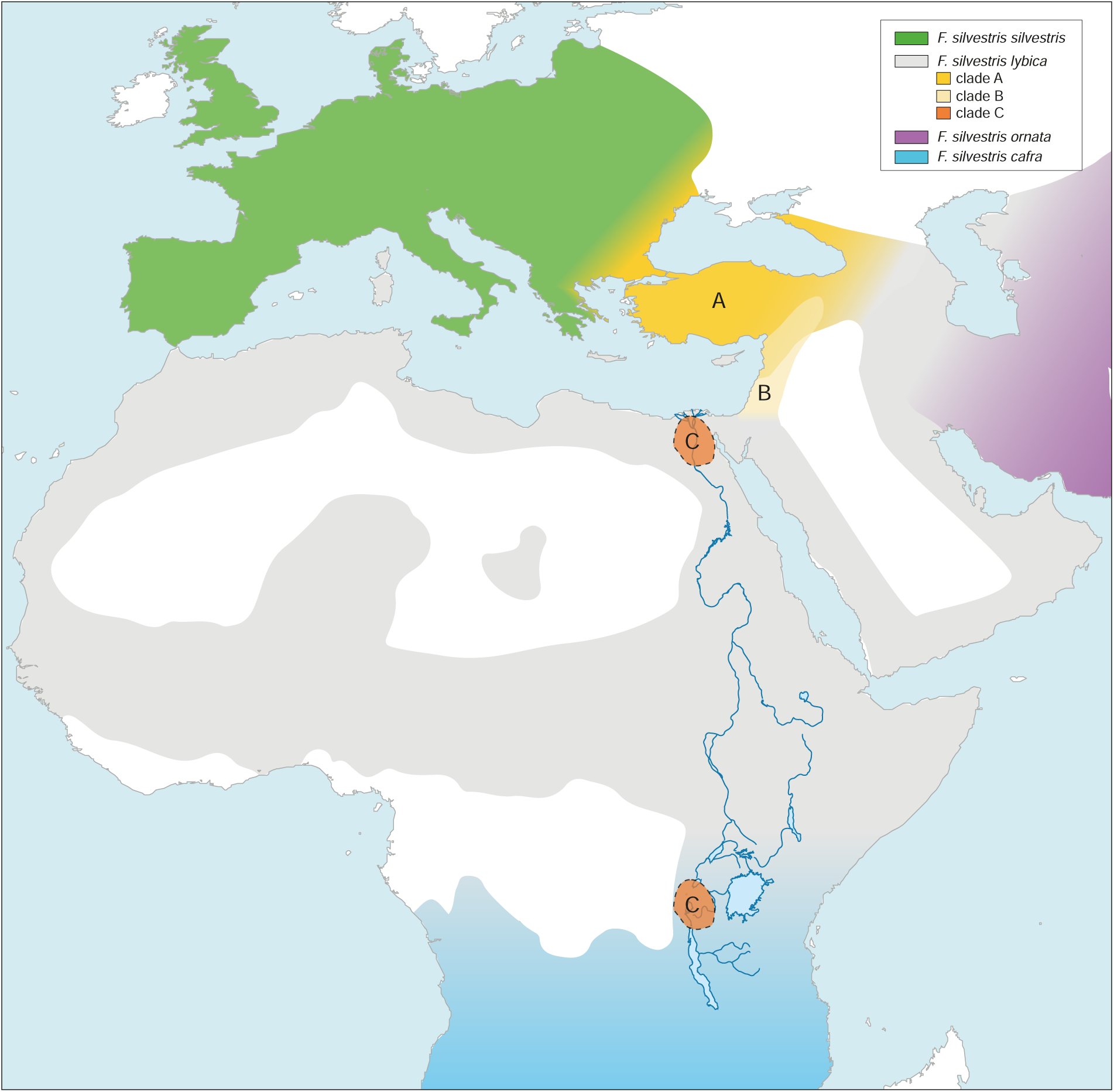
Representation of (pre)historical geographic range of *Felis silvestris* mtDNA clades in Western Eurasia and Africa as inferred in this study. The entire *F. s. lybica* (clade IV) range is in grey, and each specific clade distribution inferred from our ancient dataset has a specific color as in the tree of figure S2 and figure 1. Our dataset suggests a wide distribution of clade C on the African continent, but we reported only the areas where we directly observed it (delimited by a broken line).

